# Comparison of Gene Expression between Chronic Rhinosinusitis with Nasal Polyposis and without Nasal Polyposis

**DOI:** 10.1101/2024.11.15.623852

**Authors:** Mahmoud Fawzy Kasem, Mohamed Lotfy Helal, Ola Hamdy Elsfafey

## Abstract

**Objectives:** We aimed to compare the gene expression between chronic sinusitis with nasal polyposis and chronic rhinosinusitis without nasal polyposis either eosinophilic or not.

**Background:** Chronic rhinosinusitis (CRS) is a disease characterized by chronic inflammation of the nasal cavity and sinus mucosa. CRS mainly presents as phenotypes, namely, CRS without nasal polyps (CRSsNP) and CRS with nasal polyps (CRSwNP), based on whether it is accompanied by polyps or not. Micro-array analyses investigating the immune endo-types and pathogenesis of CRSwNP typically rely on invasive nasal biopsies.

**Methods:** This prospective randomized study was conducted on 150 patients with chronic nasal obstruction (more than 3 months). These group of patients consisted of four groups as CRSwNP patients (including eosinophilic CRSwNP [ECRSwNP] and non-eosinophilic CRSwNP [NECRSwNP]) and CRSsNP patients including eosinophilic CRSwNP [ECRSwNP] and non-eosinophilic CRSwNP [NECRSwNP]) for protein quantification by COLOC platform and gene expression evaluation by RNA-sequencing. Spearman’s analysis was performed to detect correlations between protein expression levels and clinical assessment variables.

**Results:** There were female predominance in eosinophilic CRS plus higher CT score in ECRSwNP than NECRSwNP. There was also higher SNOT-22 score in ECRSwNP than NECRSwNP. There were significant differences in all items of SNOT-22 scale after nasal polypectomy. The prevalence of the condition/disease is assumed to be 0.1. The sensitivity of 0.1 with 90% power, the specificity was 85%.

**Conclusion:** Our study revealed that there are higher recurrence, rate of manifestations for the patients with genetic predisposition of CCL18 and CCL13 more than others.

## Background

Chronic rhinosinusitis (CRS) is a disease characterized by chronic inflammation of the nasal cavity and sinus mucosa. CRS mainly presents as phenotypes, namely, CRS without nasal polyps (CRSsNP) and CRS with nasal polyps (CRSwNP), based on whether it is accompanied by polyps or not. CRSwNP patients present with severe symptoms, higher incidence of asthma, and higher recurrence rate after endoscopic nasal surgery compared with CRSsNP patients. Therefore, it is imperative to explore the pathogenesis of this chronic inflammatory disease, find key biomarkers, and elucidate the molecular basis to provide a basis for the development of new therapeutic strategies **(Wang M, et al. 2022)**.

This sub-type of CRSwNP, commonly referred to as eosinophilic CRSwNP (ECRSwNP), is characterized by nasal polyps (NPs) with strong eosinophilic infiltration and overproduction of multiple pro-inflammatory type 2 T-helper cell (Th2) -related cytokines. ECRSwNP is particularly challenging to manage as it displays rapid postoperative recurrence and is resistant to conventional treatment strategies **(Fujieda S, et al. 2019)**.

Currently, the disease is managed using steroids, which are considered the most effective treatment for achieving remission; however, their long-term use is costly and is associated with serious side effects. Thus, profiling inflammatory patterns in the nasal mucosal is crucial for understanding ECRSwNP pathogenesis and developing novel treatment strategies **(Howard BE, Lal D. 2013)**.

Micro-array analyses investigating the immune endo-types and pathogenesis of CRSwNP typically rely on invasive nasal biopsies. The common pathophysiology of the upper and lower airways has important implications for the diagnosis and treatment of respiratory co-morbidities. Previous studies on CRSwNP mainly focused on the nasal polyp tissue of the upper airway. In this study, inflamed nasal mucosal tissue and nasal polyps were used to explore deferentially expressed genes (DEGs). In addition, we combined the upper and lower airway transcriptome data of asthma patients for the analysis. DEGs identified by this approach may play a more critical role in the pathogenesis of CRSwNP **(Wang M, et al. 2022)**.

Therefore, the present study applied bio-informatics analysis to extensively explore potential genetic changes using transcriptomic data for CRSwNP and CRSsNP patients.

## Methods

This prospective randomized study was conducted on 150 patients with chronic nasal obstruction (more than 3 months). All patients with CRSwNP met the diagnostic criteria for CRSwNP according to the European Position Paper on Rhinosinusitis and Nasal Polyps 2012 guidelines. All patients with CRSsNP diagnosed with hypertrophied inferior turbinate (HIT) with chronic sinusitis more than 3 months not relived with medication. Patients were selected from general population aging from 18 up to 45 years old including patients reporting to Otorhinolaryngology outpatient clinics of our institutes.

Consent was taken from patients before performing any investigations or surgical interventions, and they had the right to refuse at any time. The study was approved by the Research Ethics Committee of the Faculty of Medicine, Menoufia University. Ethics committee reference number is 4/2021ENT/345.

Inclusion criteria were patients aged from 18 to 45 years old with CRSwNP or CRSsNP with HIT. Exclusion criteria were all patients who underwent simultaneous procedures involving the upper airway (i.e., septoplasty, inferior turbinectomy or sinus surgery) were excluded. Also, all patients with bleeding disorder were also excluded. Patients with recent upper airway infections within two weeks, choanal atresia, deviated nasal septum, and septal perforation, were also excluded using preoperative computed topography (CT) sinus. The presence of unilateral NPs, antrochoanal polyps, allergic fungal sinusitis, cystic fibrosis, or immotile cilliary disease were also exclusion criteria.

From January 2021 to January 2024, every patient data included in the study were reviewed for the following (personal data & associated symptomatology). Onset and duration of disease were also recorded. Laboratory investigations were also made for all patients preoperatively as complete blood count, bleeding profile, liver and kidney functions. Radiological assessment using Computed Topography paranasal sinuses was also reviewed to evaluate site of nasal obstruction. After that, Lund Mackay scoring system was applied to all patients’ findings in CT paranasal sinuses and opacification of osteomeatal complex (OMC). The Lund Mackay system is a widely accepted, validated staging system based on CT scan findings. The maximum score is 12 per side. Most of the studies have shown the mean Lund Mackay score in patients with CRSwNP is significantly higher than in those with CRSsNP **(Lund VJ, Kennedy DW. 1997)**.

Rhinometry and sleep lab were done to all patients to conclude nasal obstruction at time of presentation. Chest CT was also done to exclude either cystic fibrosis, or immotile cilliary disease. Following the American guidelines for allergic rhinitis, all participants were previously tested for allergies by means of the prick test then serum IgE level was done to conclude allergic consequences of hypertrophied inferior turbinates or nasal polyposis.

Preoperatively, before general anesthesia fiberoptic nasopharyngeal endoscopy was done with a 2.7 mm fiberoptic endoscope (Karl Storz, Germany) to conclude the hypertrophied inferior turbinates and to detect the presence of nasal polyposis. Either patients with CRSwNP had functional endoscopic sinus surgery or patients with CRSsNP had just partial inferior turbinectomy.

Inflamed mucosa or nasal polyp was obtained from the anterior ethmoid cavity or middle meatus. The control group comprised allergic tissues from the inferior turbinate.

The NPs or ITs specimens were fixed and processed for paraffin embedding for immune-histochemical assessment of Eosinophil inflammation. Standard 5 µm sections were prepared and stained for eosinophils (Anti-Ribonuclease 3/eosinophil cationic protein antibody, Abcam, Ab207429). The presence of ECRSwNP was determined by counting the number of eosinophils per high-power field (HPF). The histological definition of ECRSwNP was based on the presence of ≥55 eosinophils per 400 × high-power field, as previously described **(Lou H, et al. 2015)**.

We conducted COLOC analysis using the coloc package in R software (version 4.0.3). COLOC analysis assesses whether SNVs associated with gene expression and phenotype at the same locus are shared causal variants, and thus, gene expression and phenotype are “colocalized.” COLOC analysis calculates posterior probabilities (PPs) of the five hypotheses: 1) H0; no association with either gene expression or phenotype; 2) H1; association with gene expression, not with the phenotype; 3) H2; association with the phenotype, not with gene expression; 4) H3; association with gene expression and phenotype by independent SNVs; and 5) H4; association with gene expression and phenotype by shared causal SNVs. A large PP for H4 (PP.H4 above 0.75) strongly supports shared causal variants affecting both gene expression and phenotype **(Giambartolomei et al., 2014)**. We assigned a prior probability of 1 × 10^−4^ for H1 and H2 and a prior probability of 1 × 10^−5^ for H4 as the default settings of the coloc-abf function. We tested the region within 1 Mb on either side of the lead variant with the smallest *p*-value at the region in the GWAS data.

The original Sinus and Nasal Quality of Life Survey (SNOT-22) evaluates 5 clusters of symptoms (sinus infection, nasal obstruction, allergy, emotional distress, and activity limitations). Each cluster has a series of symptoms selected to help patients to understand the nature of what is being assessed. Each cluster of symptoms is rated on a 7-point Likert scale, from 0 (never) to 6 (all the time).

### Statistical Methods

Descriptive statistics included the mean value and standard deviation. The Anova and Fischer exact tests were used for the analysis of the correlation between data. The SPSS 22.0 program was used for statistical analysis. So, the p-value was considered significant as P-value less than 0.05 was considered significant.

## Results

The characteristics of the sample regarding age, sex, Lund Mackay scoring system and SNOT-22 scoring of each group are summarized in Table 1. There were female predominance in eosinophilic CRS plus higher CT score in ECRSwNP than NECRSwNP. There was also higher SNOT-22 score in ECRSwNP than NECRSwNP. SNOT-22 was also tabulated comparing between groups as shown in table 2. There were significant differences in all items of SNOT-22 scale after nasal polypectomy.

**Table 1:**
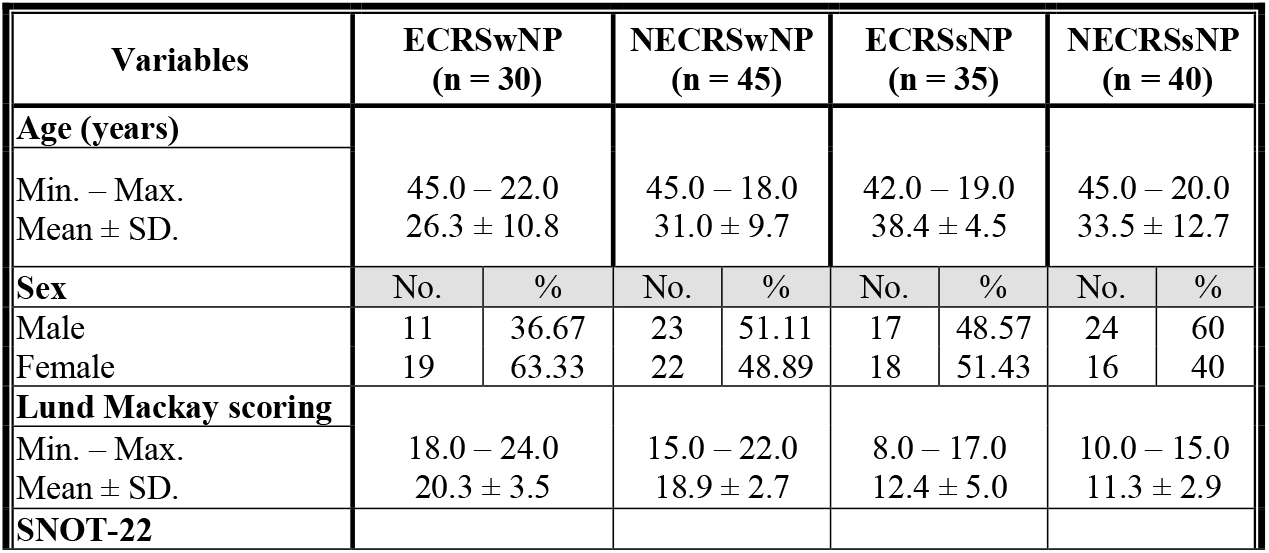
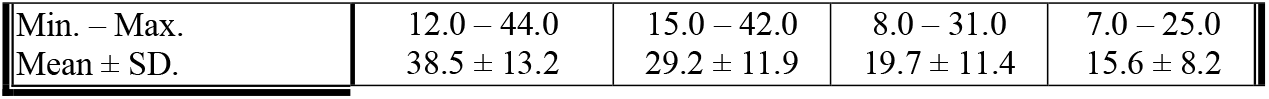
Characteristic data.

| Variables | ECRSwNP<br>(n = 30) |  | NECRSwNP<br>(n = 45) |  | ECRSsNP<br>(n = 35) |  | NECRSsNP<br>(n = 40) |  |
| --- | --- | --- | --- | --- | --- | --- | --- | --- |
| **Age (years)** |  |  |  |  |  |  |  |  |
| Min. – Max. | 45.0 – 22.0 |  | 45.0 – 18.0 |  | 42.0 – 19.0 |  | 45.0 – 20.0 |  |
| Mean ± SD. | 26.3 ± 10.8 |  | 31.0 ± 9.7 |  | 38.4 ± 4.5 |  | 33.5 ± 12.7 |  |
| **Sex** | No. | % | No. | % | No. | % | No. | % |
| Male | 11 | 36.67 | 23 | 51.11 | 17 | 48.57 | 24 | 60 |
| Female | 19 | 63.33 | 22 | 48.89 | 18 | 51.43 | 16 | 40 |
| **Lund Mackay scoring** |  |  |  |  |  |  |  |  |
| Min. – Max. | 18.0 – 24.0 |  | 15.0 – 22.0 |  | 8.0 – 17.0 |  | 10.0 – 15.0 |  |
| Mean ± SD. | 20.3 ± 3.5 |  | 18.9 ± 2.7 |  | 12.4 ± 5.0 |  | 11.3 ± 2.9 |  |
| **SNOT-22** |  |  |  |  |  |  |  |  |

|  |  |  |  |  |
| --- | --- | --- | --- | --- |
| Min. – Max. | 12.0 – 44.0 | 15.0 – 42.0 | 8.0 – 31.0 | 7.0 – 25.0 |
| Mean ± SD. | 38.5 ± 13.2 | 29.2 ± 11.9 | 19.7 ± 11.4 | 15.6 ± 8.2 |

**Table 2:** SNOT-22 scale results.

| SNOT-22 score | Preoperative | Postoperative<br>(First day) | Postoperative<br>(1 month) | P-value |
| --- | --- | --- | --- | --- |
| | Mean ± SD | | | |
| **Sinus infection** | 35.5 ± 12.0 | 18.5 ± 9.5 | 10.5 ± 3.5 | 0.035* |
| **Nasal obstruction** | 49.0 ± 15.0 | 15.5 ± 10.5 | 5.5 ± 2.5 | 0.009* |
| **Allergy** | 40.5 ± 17.0 | 21.0 ± 8.5 | 8.5 ± 3.0 | 0.043* |
| **Emotional distress** | 30.5 ± 12.0 | 20.5 ± 9.5 | 11.5 ± 7.5 | 0.025* |
| **Activity limitations** | 35.0 ± 10.0 | 19.5 ± 7.5 | 12.5 ± 5.5 | 0.021* |

\*: significant P-value, SD: Standard deviation

**Table 3:** Protein gene expression.

| Protein gene expression | ECRSwNP<br>(n = 30) | NECRSwNP<br>(n = 45) | ECRSsNP<br>(n = 35) | NECRSsNP<br>P<br>(n = 40) | Anova test |
| --- | --- | --- | --- | --- | --- |
| | Mean ± SD | | | | |
| ✓ **ANGPT1** | 8.35 ± 1.4 | 11.87 ± 1.8 | 8.96 ± 0.7 | 7.91 ± 0.9 | 0.045* |
| ✓ **CCL18** | 12.1 ± 2.1 | 11.0 ± 1.8 | 11.2 ± 1.3 | 10.5 ± 1.3 | 0.009* |
| ✓ **CCL13** | 12.9 ± 1.1 | 12.8 ± 0.9 | 12.2 ± 1.0 | 11.4 ± 1.2 | 0.013* |
| ✓ **CD40L** | 0.53 ± 0.3 | 0.41 ± 0.2 | 0.38 ± 0.3 | 0.25 ± 0.1 | 0.025* |
| ✓ **MCP4** | 13.1 ± 2.0 | 11.0 ± 2.4 | 11.6 ± 1.2 | 9.4 ± 1.1 | 0.045* |

\*: significant P-value, SD: Standard deviation

Next, we conducted the COLOC analysis to integrate the GWAS and blood eQTL data of the genes and assess whether the genes were colocalized with the trait of NP either with sinusitis or not. The COLOC test found strong support for colocalization between the trait and all the five genes (Figure 1), and thus, we considered these genes to be highly prioritized for follow-up functional studies.

**Figure 1:**
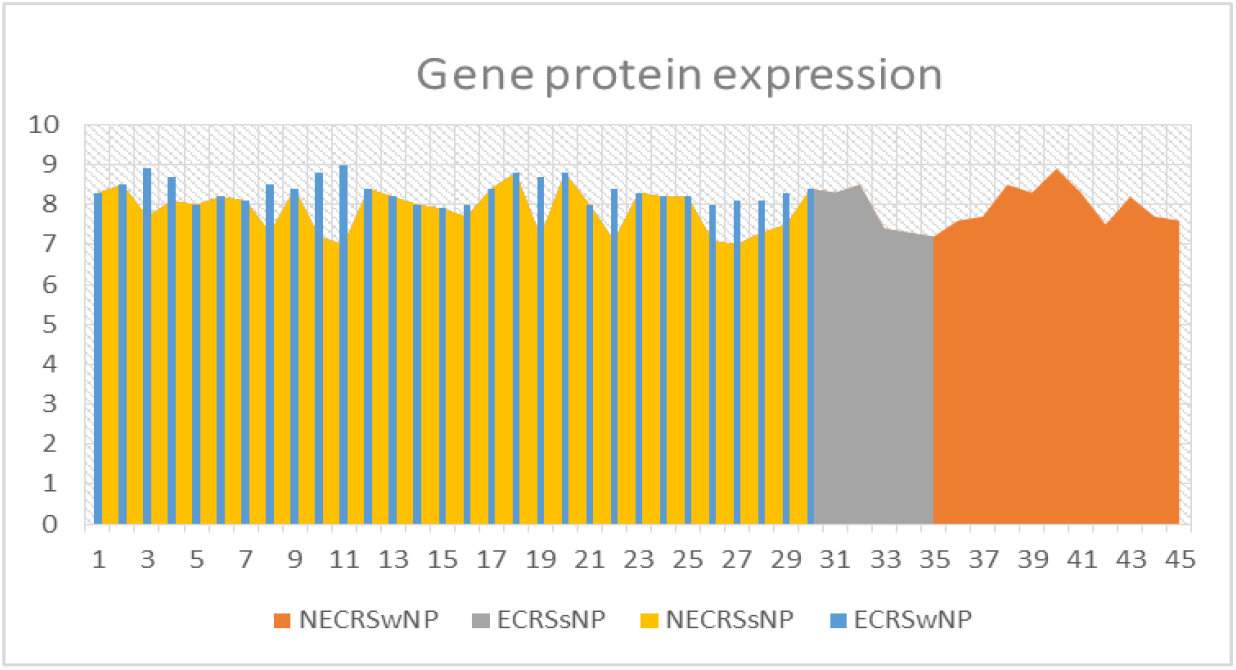
Protein gene expression coding according to quality of life.

The sensitivity of all gene expression with sinusitis recurrence comparison was made using a one-sided binomial test and the specificity comparison was be made using the same one-sided binomial test. Both comparisons were made with a Type I error rate (α) of 0.15. The prevalence of the condition/disease is assumed to be 0.1. The sensitivity of 0.1 with 90% power, the specificity was 85%.

## Discussion

COLOC analysis is one one of the most appropriate methods used for gene expression as genes related to nasal polyposis that may be genetically inherited. Integration of GWAS analysis and eQTL of blood analysis is used in this study for establishing prioritization to gene expression. There are multiple studies used the COLOC tests for detection of the expression of many genes in blood as associated with NP (TNFRSF18, CTSK, and IRF1 genes). ALOX15 and SLC22A5 genes whose associations with NP were suggested by other studies failed by COLOC tests **(Li, B., and Ritchie, M. D. 2021; Hernández C, Li X, et al. 2021)**.

Clinical and radiological manifestations could be more severe with higher scores on SNOT-22 and Lund Mackay scoring system in patients with eosinophilic CRS than non-eosinophilic as shown in our study. Many researches confirmed that eosinophilic CRS population has more aggressive clinical symptoms than other patients, but only a few studies used objective data to analyse these findings. Gitomer et al. used Lund– Mackay scores plus olfactory disability and asthma score for evaluation. Hu et al. used the VAS score to evaluate symptoms but failed to demonstrate a significant correlation between eosinophilic and non-eosinophilic CRS **(Gitomer SA, et al. 2016 ; Hu Y, et al. 2012)**.

In our study, we used the SNOT-22 score to compare preoperative values according to type of CRS, as it shows that these values were more higher in eosinophilic than non-eosinophilic and more aggressively high in eosinophilic CRS with polyposis. This could be confirmed also by higher SNOT-22 items in eosinophilic groups which further points to extra-nasal rhinological symptoms, including nasal obstruction, loss of smell, and cough. Shah et al. reviewed the pathogenesis of eosinophils accumulation that followed by degradation in eosinophilic CRS that lead to mucous secretion through release of cytokines plus chemokines. This is the explanation of this severity of these symptoms through chemotaxis **(Shah SA, et al. 2016)**.

Chemotaxis found in nasal tissues and polyps is usually associated with proteins like CCL13, CCL4, CCL19, CCL20, CCL23, MCP3, and MCP4. CCL13 usually produced in NPs could be involved in the pathogenesis of eosinophilic CRS with nasal polyps through the recruitment of multiple inflammatory cells as monocytes and macrophages. CCL13/MCP4 are potent eosinophil-modulating chemokines. They can be secreted by monocytes, macrophages that usually lead to accumulation of more eosinophils nd stimulate their degranulation. All these produce finally a cascade of inflammatory action and activation of the immune system **(Hou Y, et al. 2024)**.

Local pathology could be aggravated through production and activation of the chemokines and cytokines. These cytokines released from excess eosinophils lead to extravasation of fluid and increase the extracellular fluid, which drives eosinophils to migrate from blood stream to the vascular endothelium. Subsequently, they are recruited to the mucosal surface of the sinus lumen that lead to mucosal thickening and polyp formation **(Asano K, et al. 2020)**.

Chronic rhinosinusitis may be not only due idiopathic causes, it could be a part of genetic, or immune disorders. It could be explained as IgE-mediated allergic diseases that usually triggered by allergen materials that release cytokines from basophils. A majority of European cases with rhinosinusitis with nasal polyps are usually characterized by type 2 inflammation with eosinophilia and elevated levels of type II cytokines such as interleukin-5 and 13 **(Hulse et al., 2015; Hopkins, 2019)**.

Although a lot of studies made on different genes expression and their effects as causal events for nasal polyps through eosinophilia. Our analysis was made on ANGPT-1 and CD40L had prioritization in gene expression on nasal polyposis. Further studies would be required to examine the effect of ANGPT-1 gene on mucosal changes of CRS with NP.

This study had several major limitations. First, we may have missed some important genes that have functional associations with NP, especially because of the small sample size of the GWAS on NP. Second, our COLOC analyses were based on populations predominantly from the middle east. Therefore, our findings are unlikely to be generalized to all populations.

## Conclusion

In conclusion, we could prioritize several genes associated with NP, including CCL18, CCL13, and CD40L, by COLOC analyses using the latest GWAS data with 34 genome-wide significant loci and blood eQTL data. However, follow-up functional studies must be conducted in future to validate the functional associations of the genes with the underlying disease mechanisms.

## Acknowledgments

Not applicable.

